# Efflux Pump Activation Confers Mupirocin Resistance and Enhances Rhizosphere Fitness in *Pseudomonas*

**DOI:** 10.64898/2026.01.05.697689

**Authors:** Wen-Jie Wang, Man-Jin Wang, Wen-Jun Jiang, Li-Qun Zhang

## Abstract

Mupirocin (Mup) is a polyketide antibiotic for clinical MRSA treatment and agricultural biocontrol, acting by binding to IleRS to inhibit protein synthesis. Here, we demonstrate that *Pseudomonas viciae* 11K1 acquires resistance to mupirocin not through canonical mutations in *ileS*, the gene encoding the drug target, but via single nucleotide polymorphisms (SNPs) of EmhR, a transcriptional repressor. These SNPs attenuate EmhR’s DNA-binding ability, resulting in derepression of the resistance-nodulation-division (RND) efflux pump EmhABC. This leads to a 7-fold (to 800 μg/mL) increase in the mupirocin minimum inhibitory concentration (MIC), and confers cross-resistance to multiple other antibiotics. Critically, 11K1 strains harboring EmhR mutations exhibit enhanced competitive fitness in colonizing wheat rhizospheres against the mupirocin-producing *Pseudomonas bijieensis* 2P24. The conservation of this regulatory mechanism in *Pseudomonas aeruginosa*, mediated by the EmhR ortholog NalD, underscores its broader biological significance. Our findings establish a direct link between efflux pump regulation and ecological adaptation, highlighting a key mechanism contributing to environmental antimicrobial resistance with important implications for the clinical and agricultural use of mupirocin.

**IMPORTANCE:** Mupirocin and its producing *Pseudomonas* strains are widely used in both clinical and agricultural contexts, making mupirocin resistance a significant concern for public health and food security. While canonical mupirocin resistance is primarily attributed to mutations in *ileS*, the gene encoding the drug target isoleucyl-tRNA synthetase (IleRS), our study identifies a novel resistance mechanism in *Pseudomonas viciae* strain 11K1 mediated by SNPs in the transcriptional repressor EmhR. These SNPs derepress the RND efflux system EmhABC, and reduce intracellular mupirocin levels. This mechanism not only enhances tolerance to mupirocin but also confers cross-resistance to multiple antimicrobial agents, raising the risk of multidrug-resistant strain spread. EmhR-mutant 11K1 strains exhibit enhanced rhizosphere competitiveness in the presence of mupirocin-producing *Pseudomonas bijieensis* strain 2P24, indicating an ecological fitness advantage that could promote resistant population expansion in agriculture environments. The same regulatory pathway is conserved in human pathogen *Pseudomonas aeruginosa* PAO1 via the functional ortholog NalD, suggesting that this mechanism may be widespread in *Pseudomonas*. These findings fill a critical gap in understanding non-target-based mupirocin resistance, clarify the ecological drivers of antimicrobial resistance (AMR), and offer practical insight for improving mupirocin application to limit the emergence and spread of resistant pathogens.

## INTRODUCTION

Antimicrobial resistance (AMR) represents a critical threat to global public health and ecological security. The widespread dissemination of multidrug-resistant (MDR) bacteria across medical systems, food chains, and natural environment has led to substantial increases in healthcare expenditures and environmental management costs (1–3). Antibiotic resistance is fundamentally an adaptive evolutionary response of microorganisms to the selective pressure of antibiotics. Established resistance mechanisms include: i) production of enzymes that inactivate antibiotics; ii) mutations in target sites that reduce antibiotic binding affinity; iii) active efflux of antibiotics via efflux pump systems, particularly the resistance nodulation cell-division (RND) family; and iv) reduced drug permeability due to structural features of the cell membrane (4–7). In soil ecosystems, although the rate of antibiotic resistance emergence is lower than in medical environments, the robust colonization potential of antibiotic-producing bacteria may impose selective pressure on cohabiting indigenous microbial populations, potentially driving the acquisition of resistance through adaptive evolution. Elucidating the evolutionary mechanisms of microbial adaptation to antibiotics is crucial for understanding competitive interactions and the development of antibiotic resistance within the rhizosphere ecosystem.

Mupirocin, also known as pseudomonic acid A, is a polyketide antibiotic widely utilized in the treatment of skin and nasal infections (8, 9). It was originally identified in bacteria belonging to the genus *Pseudomonas*, which are prevalent members of the rhizosphere microbiota and frequently used for the biological control of soil-borne plant diseases (10–14). mupirocin primarily targets isoleucine-tRNA synthetase (IleRS), an essential enzyme in protein translation. Its epoxide-containing side chain structurally mimics Ile-AMP, allowing it to competitively inhibit IleRS by blocking the incorporation of isoleucine into nascent polypeptide chains (15–17). mupirocin-producing *Pseudomonas* strains may exploit this compound to gain a competitive advantage in the rhizosphere (18). Meanwhile, coexisting bacterial population must develop adaptive mechanisms to withstand mupirocin exposure to ensure survival. mupirocin resistance commonly arises through expression of IleRS variants with reduced affinity, such as eukaryotic IleRS or prokaryotic IleRS2 homologs (19–21), or via point mutations in the *ileS* gene that diminish mupirocin binding (22–24). In Gram-negative bacteria, resistance is further mediated by resistance-nodulation-division (RND) efflux pump systems, tripartite complexes that span the inner membrane, periplasm, and outer membrane (OM). Examples include AcrAB-TolC in *E. coli* and MexAB-OprM in *P. aeruginosa* (25–27). Notably, *E. coli tolC* deletion mutants exhibit markedly increased susceptibility to mupirocin (28), indicating that the efflux activity contributes significantly to microbial tolerance of this antibiotic.

In the plant rhizosphere ecosystem, mupirocin-producing *Pseudomonas* strains may gain a competitive advantage by inhibiting surrounding microbial competitors through antibiotic activity. However, understanding of how how mupirocin-sensitive rhizobacteria adapt to this selective pressure remains limited. In this study, using the mupirocin-producing strain *Pseudomonas bijieensis* 2P24 and mupirocin-sensitive strain *Pseudomonas viciae* 11K1 as model systems, we demonstrated that exposure to mupirocin induces point mutations in the transcriptional regulator EmhR in 11K1. These mutations result in the upregulation of the RND-type efflux pump EmhABC, thereby enhancing mupirocin resistance. The increased expression of the EmhABC significantly improves the competitive colonization ability of 11K1 in wheat rhizosphere in the presence of 2P24. Our findings indicate that the regulatory modulation of efflux pump expression constitutes a key mechanism underlying bacterial ecological adaptation to antibiotic stress.

## RESULTS

### *Pseudomonas viciae* 11K1 is sensitive to mupirocin produced by *P. bijieensis* 2P24

Both *Pseudomonas* strains 2P24 and 11K1 were isolated from the plant rhizosphere. 2P24 produces polyketide antibiotics mupirocin and Diacetylphloroglucinol (DAPG) (Figure 1C) (29), whereas 11K1 produces lipopeptide antibiotics brasmycin and braspeptin (30). On PDA plates with glucose as the carbon source, strain 2P24 was observed to significantly inhibit the growth of 11K1 (Figure 1A). Analysis using 2P24 mutants, specifically the mupirocin-deficient mutant (Δ*mup*), DAPG-deficient mutant (Δ*phlD*), and the double mutant (Δ*phlD*Δ*mup*), revealed that Δ*phlD* exhibited inhibitory activity against 11K1 comparable to that of the wild-type 2P24, whereas Δ*mup* and Δ*phlD*Δ*mup* showed markedly reduced inhibition (Figure 1A). These findings indicate that mupirocin is the key antibiotic responsible for 2P24-mediated inhibition of 11K1. Consistent results were obtained in the liquid LB medium using purified compounds: mupirocin inhibited the growth of 11K1 in a concentration-dependent manner, with a minimum inhibitory concentration (MIC) of 0.3 mM (100 μg/mL), while DAPG exhibited no inhibitory effect on 11K1 across the tested concentration range of 7.5-500 μM (Figure 1B). Collectively, these results demonstrate that mupirocin is the key antimicrobial agent underlying the suppression of 11K1 by 2P24.

**Figure 1.**
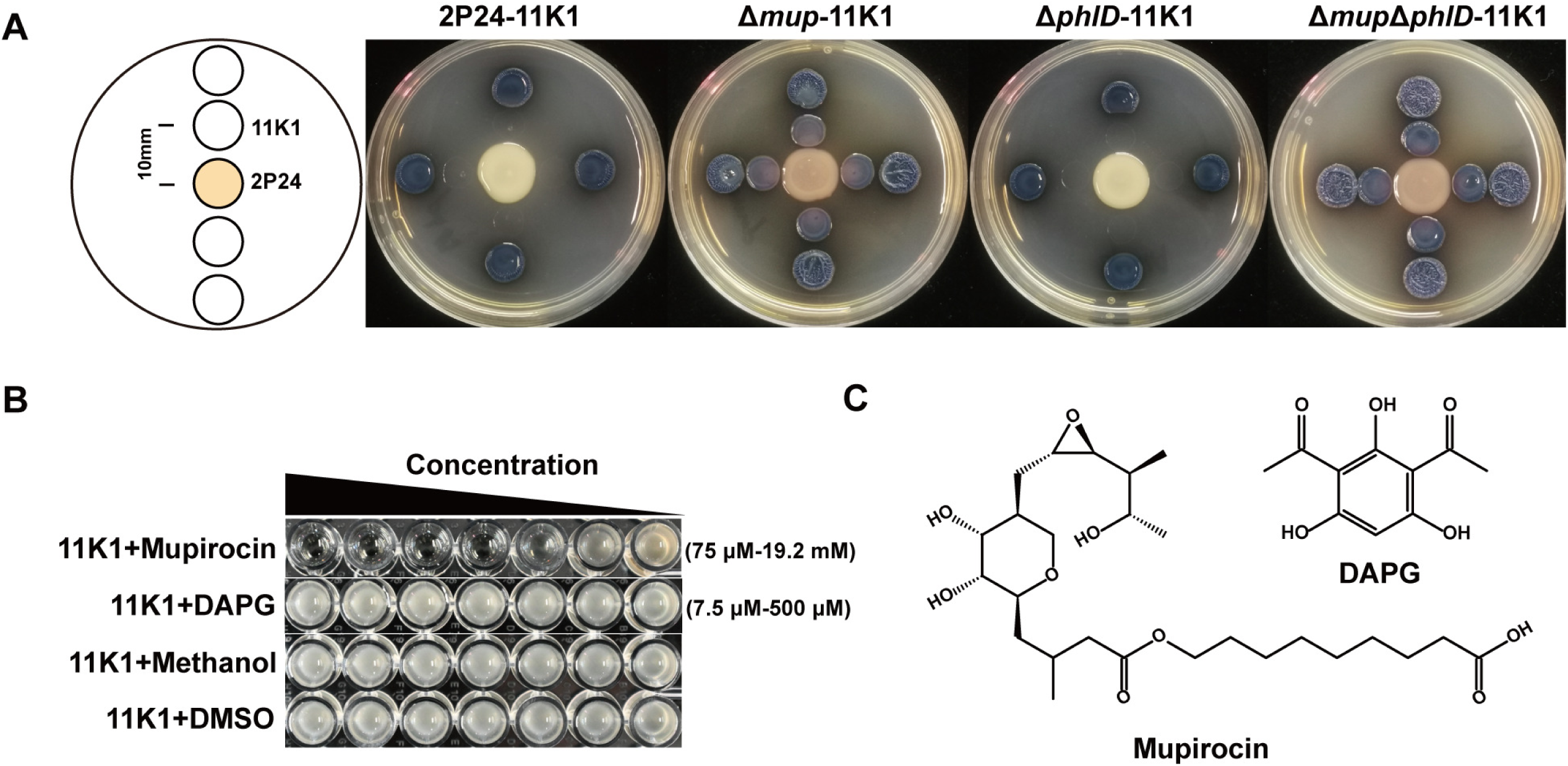
Identification of the key antibiotic by 2P24 that inhibits 11K1. A. A 5-μL drop of 2P24 suspension or its antibiotic-deficient mutant was spotted on the center of the PDA agar plates and incubated at 28°C for 24 hours. Wild-type strain 11K1 was then inoculated at distances of 10 mm and 20 mm from the 2P24 colonies. Inhibition zones were observed after an additional 24 h of incubation at 28°C. B. Determination of the minimum inhibitory concentrations (MICs) of mupirocin and 2,4-diacetylphloroglucinol (DAPG) against 11K1. C. Chemical structures of mupirocin and DAPG.

### Development of mupirocin-resistance mutants in strain 11K1 under selective pressure

To generate mupirocin-resistant mutants of strain 11K1, 10 μL of 10 mg/mL mupirocin was applied to the center of a King’s B agar plate freshly inoculated with 11K1. After incubation of 18 hours, mupirocin diffusion resulted in a clear inhibition zone. Colonies appearing at the edge of the inhibition halo, where the antibiotic concentration is relatively low, were observed (Figure 2A). These colonies were considered potential mupirocin-resistant variants and were purified for subsequent MIC assays (Supplementary Figure S2A). Five isolates were identified as mupirocin-resistant mutants, with the MIC values increased four to eight fold compared to their parental strain 11K1 (defined as the starting wild-type strain in the experiment) (Figure 2B).

**Figure 2.**
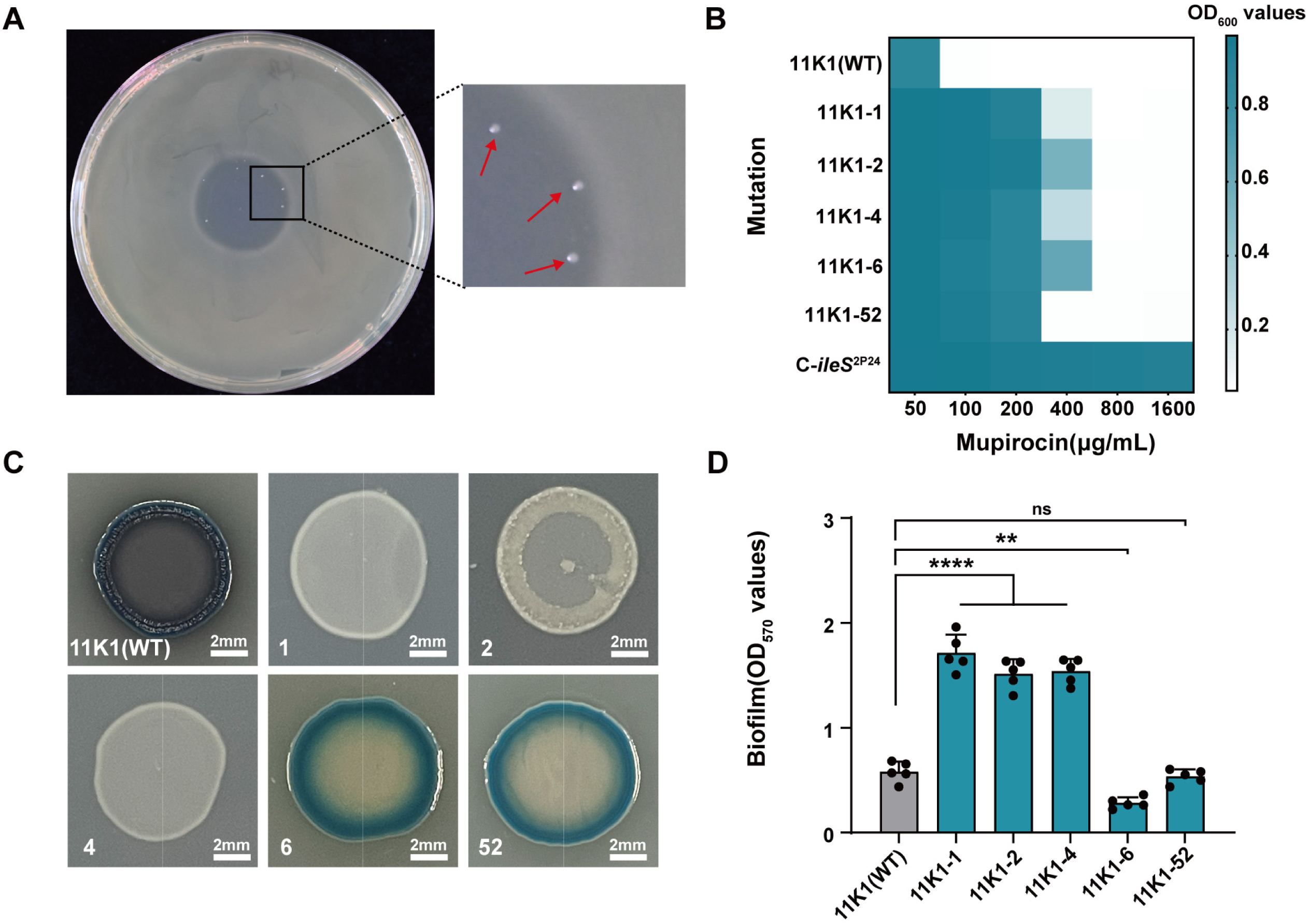
Generation and screening of mupirocin-resistant mutants of strain 11K1. A. 10-μL aliquot of 10 mg/mL mupirocin was applied to the center of a King’s B agar plate inoculated with wild-type 11K1. Colonies growing at the interface between the mupirocin inhibition zone and the bacterial lawn (indicated by red arrows) were selected as candidate mupirocin-resistant mutants. B. Heatmap displaying mupirocin MIC values for 11K1 and its derived mutants: The x-axis represents mupirocin concentrations (μg/mL), and the left y-axis lists the parental strain 11K1 and its derivative mutants. The entry “C*-iles*^2P24^” refers to the complemented strain in which the *ileS* gene, a mupirocin tolerance determinant from strain 2P24 exhibiting low drug affinity, was introduced into the parental 11K1 background. The right y-axis indicates the optical density at 600 nm (OD₆₀₀), reflecting bacterial growth levels. C. Phenotypic characteristics of 11K1 and mupirocin-resistant mutants grown on PDA plates. Scale bars = 2 mm. D. Biofilm formation capacity of 11K1 and mupirocin-resistant mutants, quantified by OD₅₇₀ measurements after crystal violet staining. Statistical significance was determined by one-way analysis of variance (ANOVA) with Tukey’s post-hoc test: **P < 0.01, ****P < 0.0001; ns, not significant.

Although no significant differences in growth were observed for the mupirocin-resistant mutants compared to the parent strain 11K1 in LB or M9 liquid medium (Supplementary Figure S2B and S2C), distinct phenotypic changes were observed in colony level. Strain 11K1 produced an indigo pigment on PDA solid medium, while the mutants 11K1-1, 11K1-2, and 11K1-4 lost this ability, forming milky white colonies with reduced surface wrinkling (Figure 2C, Supplementary Figure S2D). Additionally, the biofilm formation was significantly enhanced in the mutants 11K1-1, 11K1-2, and 11K1-4 relative to the parental strain. In contrast, mutant11K1-6 exhibited markedly reduced biofilm production, while mutant 11K1-52 showed no significant difference compared to 11K1(Figure 2D).

### Genome sequencing and SNP analysis of the mupirocin-resistant mutants of strain 11K1

Although mupirocin-resistant mutants of 11K1 exhibited increased resistance compared to the wild-type strain, their resistance levels remained substantially lower than that of the 11K1 expressing the exogenous *ileS*^2P24^ gene, indicating that the spontaneous mutations in 11K1 did not fully confer high-level resistance to mupirocin (Figure 2B). In methicillin-resistant *Staphylococcus aureus* (MRSA) and *Escherichia coli* MC4100, single-nucleotide mutations in the *ileS* gene (e.g., V588F, V631F and G593V) have been shown to enhance bacterial resistance to mupirocin (22, 31, 32). We therefore performed sequencing analysis of the *ileS* gene amplified by PCR from these resistant 11K1 mutants (Supplementary Table S2), but no mutations were detected (data not shown).

We further conducted whole-genome re-sequencing of the parent 11K1 and five mutant strains using the Illumina MiSeq platform. Sequencing data are analyzed for single-nucleotide polymorphisms (SNPs), and candidate variants were validated by PCR amplification and Sanger using gene-specific primers (see Supplementary Table S2). A total of five genetic variants were identified across the five mutants, predominantly consisting of nonsynonymous amino acid mutations and frameshift mutations caused by nucleotide insertions (Figure 3A). The majority of these mutations occurred in the response regulator gene *gacA* of the GacS/GacA two-component system and in *emhR*, which encodes a TetR family transcriptional regulator. In *gacA*, insertion of a guanine (G) after position 372 results in a frameshift and premature truncation of the protein. This mutation is present in the mutants 11K1-1, 11K1-2, and 11K1-4. Additionally, mutant 11K1-52 carries a non-synonymous substitution in *gacA* (L80R). Mutations in *emhR* were identified in strains 11K1-1 and 11K1-6, both harboring the A47P substitution. Strain 11K1-6 also contains an additional I112T substitution in *emhR*, whereas 11K1-1 has a frameshift-causing insertion at position 295 of the *emhR* coding sequence. In other mutants, 11K1-2 carries nonsynonymous mutations in *soxR*, encoding a MerR family transcriptional activator, and 11K1-52 harbors a mutation in *secDF*, which encodes a heterodimeric membrane complex involved in inner membrane protein transport.

**Figure 3.**
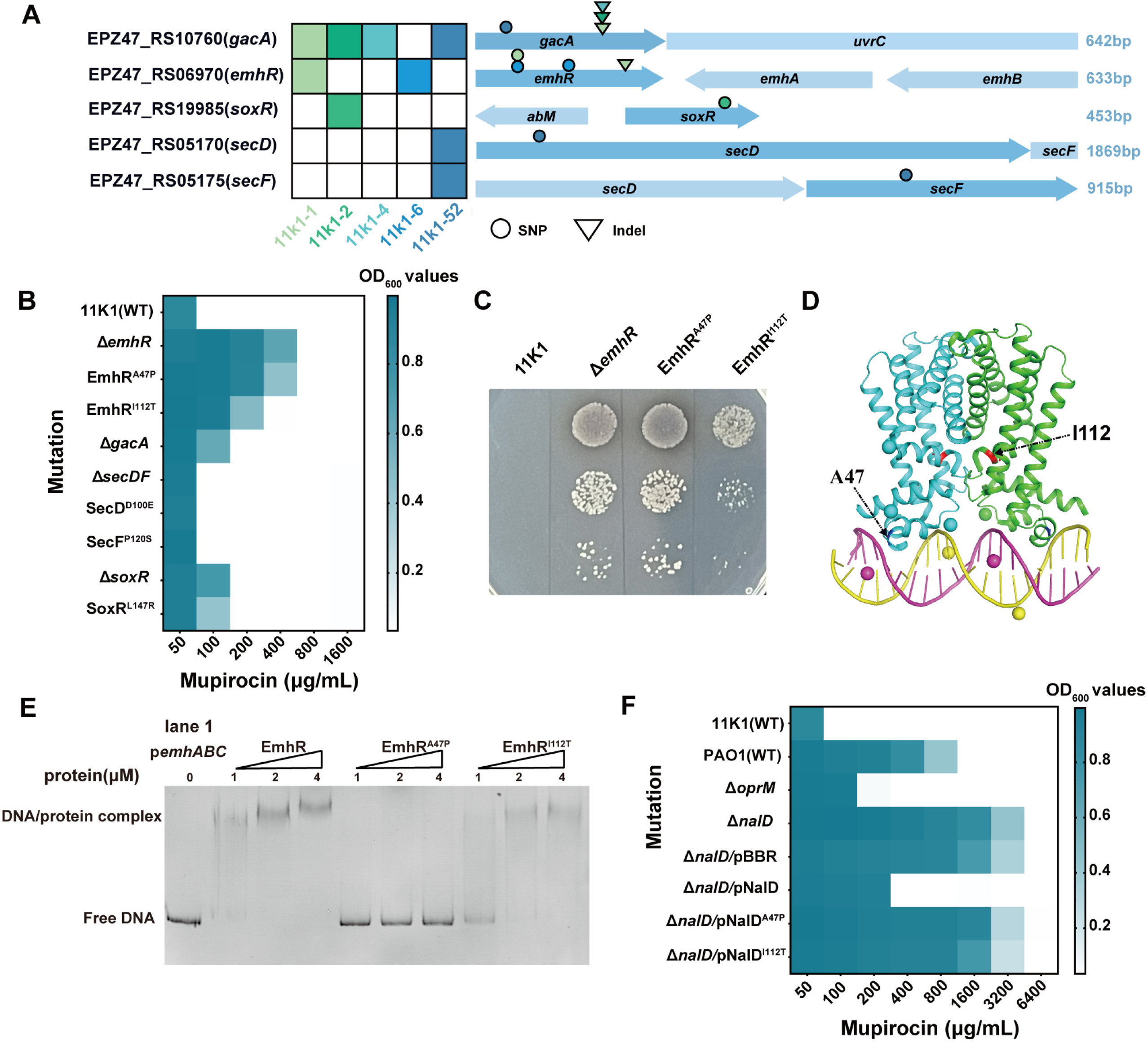
EmhR is a key regulator mediating mupirocin resistance in strain 11K1. A. Heatmap of mutated genes and associated mutation types in mupirocin-resistant 11K1 mutants. Left: Locus tags (e.g., EPZ47_RS10760) and gene names (e.g., *gacA*) of target genes. Middle: Schematic representation of open reading frames (ORFs) of target genes (dark blue) and adjacent genes (light blue). Right: Nucleotide lengths of the target genes. Mutation types are indicated by symbols: spheres represent nonsynonymous amino acid mutations; inverted triangles denete nucleotide insertions. B. mupirocin MIC assays comparing strain 11K1 with its derived mutants harboring single-nucleotide polymorphisms (SNPs) or deletions in candidate genes, constructed via homologous recombination. C. Sensitivity of 11K1, the *emhR* deletion mutant (Δ*emhR*), and EmhR point mutants (EmhR^A47P^, EmhR^I112T^) to mupirocin (100 μg/mL) on solid PDA plates. D. Homology model of the EmhR dimer bound to its target DNA, predicted by AlphaFold 3 based on the MtrR crystal structure as a template, and visualized with PyMOL. Positions of amino acid residues A47 and I112 are indicated by black arrows. E. Electrophoretic mobility shift assay (EMSA) was used to assess the interaction between purified His6-EmhR (1-4 μM) and the *emhABC* promoter region (p*emhABC*), with the buffer-only sample serving as the negative control (lane 1). F. MIC assays of mupirocin against *Pseudomonas aeruginosa* PAO1 and its derivative mutants. The construct Δ*nalD*/pNalD refers to the Δ*nalD* mutant complemented with the *nalD* fragment carried by the plasmid pBBR1MCS-5; other constructs follow the same nomenclature.

### EmhR is a critical determinant underlying enhanced mupirocin resistance in strain 11K1

To identify the key sites responsible for mupirocin resistance in the 11K1 mutants, we constructed single-gene deletion mutants of *gacA*, *soxR*, *secDF*, and *emhR*, as well as site-directed single-nucleotide mutants at identified SNP loci within these genes. MIC assays showed that, compared to the parental strain 11K1 (MIC: 100 μg/mL), the Δ*gacA* and Δ*soxR* mutants exhibited a twofold decrease in resistance (MIC: 200 μg/mL), while the Δ*emhR* mutant showed an eightfold reduction in susceptibility (MIC: 800 μg/mL). These results indicate thatEmhR plays a more critical role in the mupirocin resistance than GacA and SoxR. Analysis of single-base substitution variants further supports this conclusion: both EmhR^A47P^ and EmhR^I112T^ significantly enhance resistance to mupirocin, with MIC values 8-fold (800 μg/mL) and 4-fold (400 μg/mL) higher than that of the wild type, respectively (Figure 3B). In contrast, the SoxR^L147R^ mutant displayed resistance levels comparable to those of the Δ*soxR* and Δ*gacA* mutants. Moreover, neither the point mutants SecD^D100E^ and SecF^P120S^ nor the *secDF* deletion mutant showed altered MIC values relative to the parental strain. These findings suggest that the heterodimeric inner membrane protein transporter SecDF has minimal involvement in modulating mupirocin resistance in strain 11K1. In summary, MIC assay data demonstrate that EmhR is a crucial regulatory factor governing mupirocin resistance in strain 11K1. Notably, the two EmhR SNP variants confer distinct degrees of resistance enhancement, with the alanine residue at position 47 appearing to play a more prominent role (Figure 3B and 3C).

In *Pseudomonas* species, EmhR functions as a transcriptional regulator that specifically recognizes and binds to a conserved 40bp characteristic DNA sequence (33), thereby negatively regulating the expression of the RND-type efflux pump EmhABC. This regulatory mechanism promotes the extrusion of various toxic compounds from the cells (34). Leveraging the structural information the EmhR homolog MtrR (PDB code 3VIB), we used AlphaFold 3 and PyMOL to predict the interaction model between the EmhR and its target DNA (Figure 3D). Structural analysis revealed that residue A47 of EmhRis located within the α3 helix of the TetR-family regulator and is in close proximity to the DNA-binding interface of the EmhR dimer, whereas I112 resides near a critical region involved in the dimerization between EmhR monomers. The EMSA further elucidated the functional significance of these two SNP sites. EmhR bound specifically to the promoter region of *emhABC* in a concentration-dependent manner (Figure 3E). Competitive binding assays using FAM-labeled *emhABC* promoter DNA (FAM-p*emhABC*) and unlabeled p*emhABC* confirmed the specificity of EmhR–DNA interactions (Supplementary Figure S3). Both EmhR^A47P^ and EmhR^I112T^ variants exhibited markedly reduced DNA-binding capacities, with the A47P substitution showing a more pronounced effect, nearly abolishing DNA binding (Figure 3E). These findings suggest that under mupirocin selective pressure, strain 11K1 acquires point mutations that impair EmhR stability or its DNA-binding affinity, leading to derepression of the EmhABC efflux system and consequently enhanced antibiotic resistance.

To further validate the functional conservation of the EmhR-like mutations in enhancing bacterial resistance to mupirocin, we identified NalD, the ortholog of EmhR in 11K1, in the human opportunistic pathogen *P. aeruginosa* PAO1. Unlike MexR, the primary transcriptional regulator that binds to the promoter region of MexAB-OprM,, NalD directly represses the efflux gene expression of by binding to a second upstream promoter element of the MexAB-OprM system (35). MIC assays revealed that, compared to the wild-type PAO1 (MIC:1600 μg/mL), the Δ*oprM* mutant exhibited a sevenfold decrease in resistance (MIC: 200 μg/mL) (Figure 3F), indicating a strong association between the RND-type efflux pump MexAB-OprM and mupirocin adaptation. Conversely, the Δ*nalD* mutant displayed a threefold increase in resistance (6400 μg/mL). Notably, complementation of the Δ*nalD* mutant with the intact *nalD* gene restored the susceptibility, reducing the MIC value threefold (MIC: 400 μg/mL). However, complementation with either NalD^A47P^ or NalD^I112T^ failed to restore repression, resulting in mupirocin resistance levels comparable to those of the Δ*nalD* mutant. These findings not only indicate that NalD-mediatednegative regulation of MexAB-OprM significantly influences bacterial adaptation to mupirocin, but also suggest that residues A47 and I112 are critical for the regulatory function of NalD.

### EmhR-Mediated Upregulation of EmhABC expands the Antibiotic Resistance profile of strain 11K1

EmhR has been identified as a negative regulator of the transcription of the RND-type efflux pump genes *emhABC* in *Pseudomonas* spp.(36). The point mutations A47P and I112T, which induce structural alterations in EmhR, are likely to affect *emhABC* expression. Reverse transcription quantitative PCR (RT-qPCR) assays revealed that, compared to the wild-type strain 11K1, the transcriptional levels of *emhA*, *emhB*, and *emhC* were significantly upregulated in the Δ*emhR* mutant at both 12 and 18 hours post cultivation (Figure 4A and Supplementary Figure S3). These findings confirme the repressive role of EmhR in *emhABC* transcription and are consistent with previous studies (33, 37). Notably, the EmhR^A47P^ mutant exhibited a regulatory effect comparable to that of the Δ*emhR* deletion mutant, with both strains showing marked upregulation of *emhABC* transcripts. In contrast, the EmhR^I112T^ mutation displayed a weaker effect, significantly enhancing only *emhA* transcription at the 12-hour time point. This observation is consistent with the EMSA data, which demonstrated that the I112T variant retains a higher DNA-binding affinity to the target sequence than the A47P mutant. MIC assays indicated that the Δ*emhABC* mutant exhibited a 16-fold reduction in mupirocin resistance compared to the wild type, highlighting the essential role of the EmhABC efflux pump in conferring mupirocin resistance. Moreover, both the *emhR* deletion mutation and the A47P and I112T point mutations displayed 4 to 8-fold increased resistance to mupirocin relative to the wild-type strain 11K1, likely due to derepression of EmhABC resulting from the impaired EmhR function. Collectively, these data highlight the critical contribution of EmhR mutations to enhanced *emhABC* expression and acquired mupirocin resistance in strain11K1.

**Figure 4.**
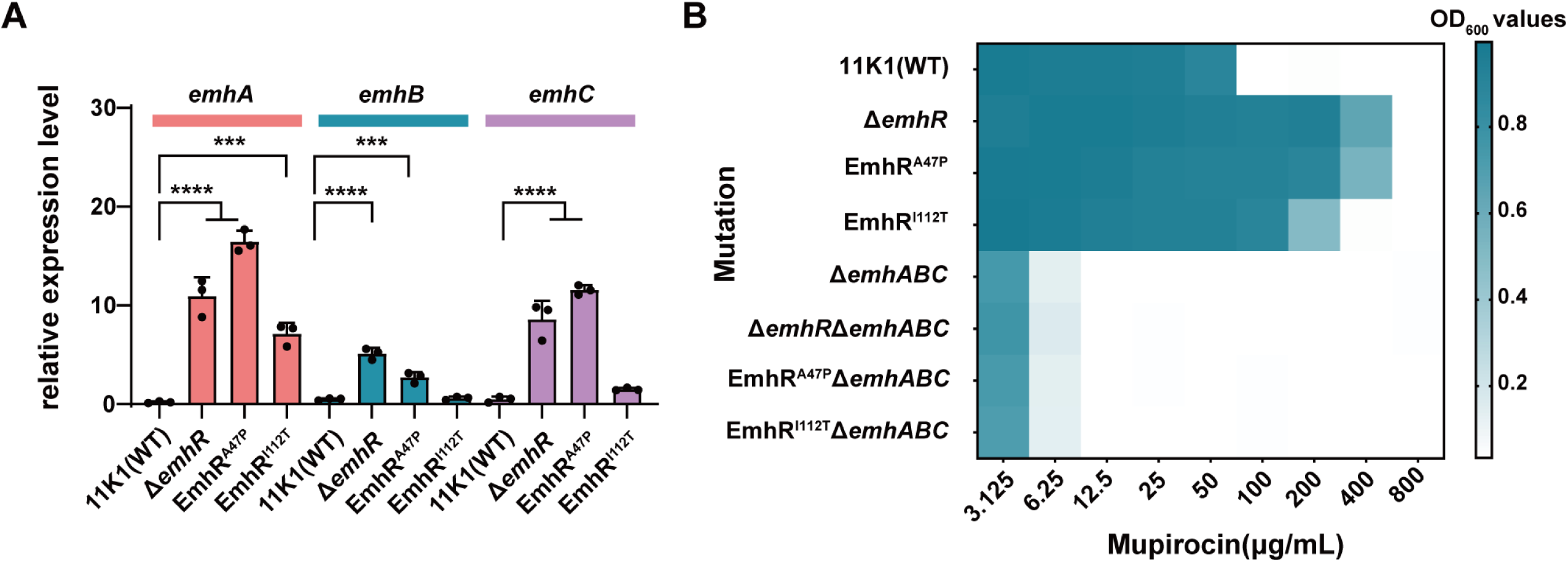
EmhR deletion and point mutations regulate *emhABC* transcription and mupirocin resistance. A. Transcriptional expression levels of *emhA*, *emhB*, and *emhC* in 11K1, the Δ*emh*R mutant, and EmhR point mutant (EmhR^A47P^, EmhR^I112T^) at 12 hours post-inoculation (hpi), quantified by quantitative reverse transcription PCR (qRT-PCR). B. MIC heatmap of mupirocin against 11K1 and its derived mutants. The x-axis represents mupirocin concentrations (μg/mL), the left y-axis lists bacterial strains, and color intensity indicates OD₆₀₀ values. Statistical significance was determined by one-way ANOVA with Tukey’s post-hoc test: ****P < 0.0001.

The upregulated expression of EmhABC generally enhances the efflux of various antibiotics and cytotoxins in bacteria. Therefore, we performed MIC assays using multiple bacteriostatic agents to evaluate the resistance profiles of EmhR mutant. The results showed that the deletion of the entire *emhABC* gene cluster significantly reduced resistance to multiple antibiotics, suggesting that EmhABC functions as a crucial broad-spectrum resistance determinant in strain 11K1. Moreover, point mutations in EmhR, particularly the A47P substitution, not only increased the resistance to mupirocin but also significantly enhanced resistance to ampicillin, gentamicin, tetracycline, and chloramphenicol (Table 1), with an effect comparable to that of the Δ*EmhR* mutant. These findings suggest that some key point mutation in EmhR, such as EmhR^A47P^, expand the antibiotic resistance spectrum and confer enhanced environmental adaptability and survival advantages in complex and variable ecological settings, particularly under antimicrobial selective pressure. In contrast, resistance levels to certain agents, such as Polymyxin B, were unaffected by mutations in the EmhR-EmhABC efflux pump, likely because the mechanism of action of Polymyxin B is independent of this efflux pathway (38).

**Table 1.**
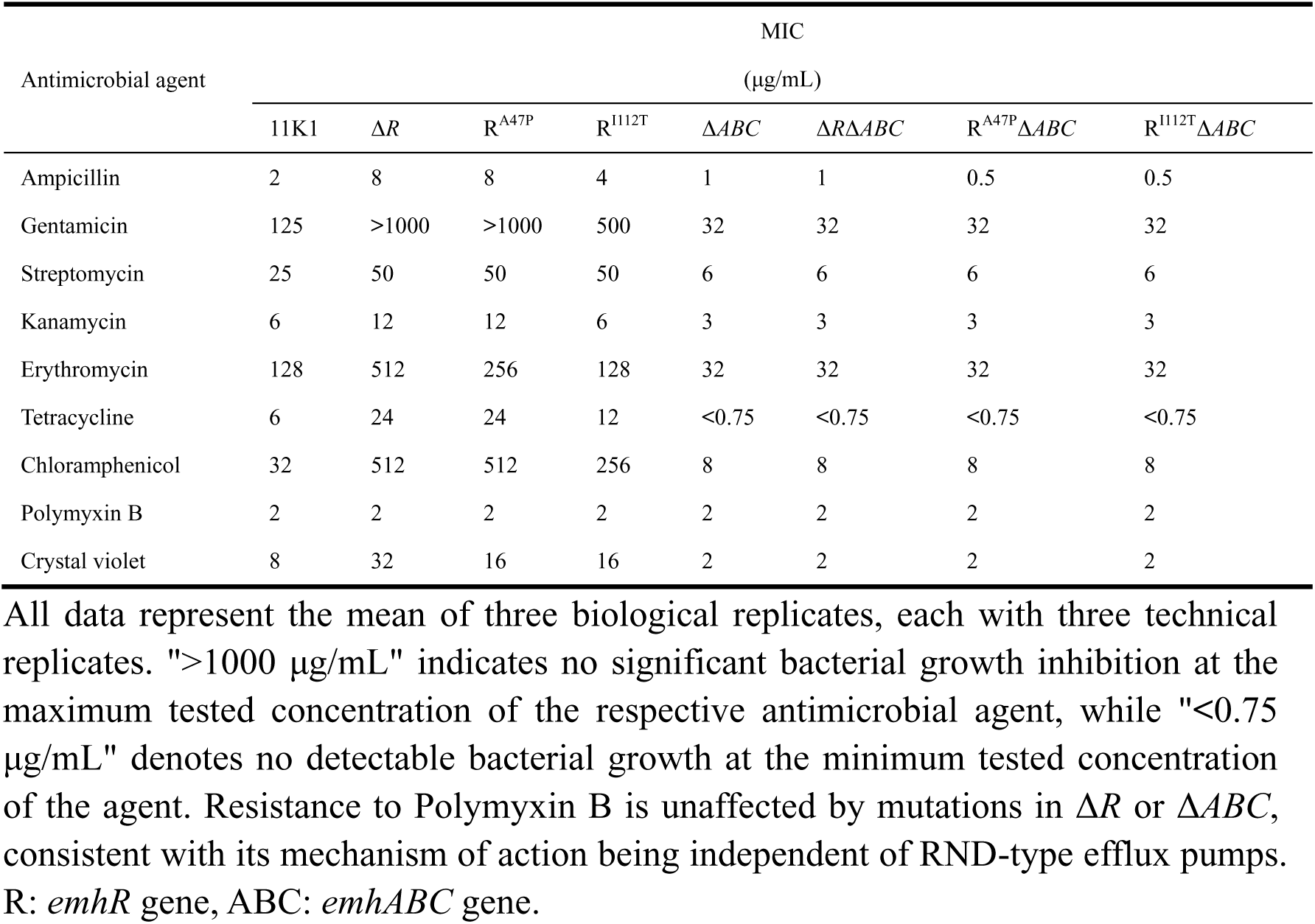
MICs of various antimicrobial agents against 11K1 and its derived mutants.

### Point mutations in EmhR enhance the competitive colonization ability of strain 11K1 in the rhizosphere

Antimicrobial resistance is a crucial mechanism enabling bacterial adaptation to ecological environments. To evaluate the impact of A47P and I112T point mutations in EmhR on rhizosphere colonization capacity of strain 11K1, competitive colonization assays were conducted against strain 2P24. The results revealed that, compared to the wild-type strain 11K1, the Δ*emhABC* mutant exhibited significantly reduced colonization in the wheat rhizosphere (Figure 5A), indicating the essential role of the EmhABC efflux pump in the niche competition within this environment. Notably, statistical analysis of the absolute colonization levels in sterilized soil demonstrated that while the EmhR point mutations (A47P and I112T) and the Δ*emhR* mutant did not alter the intrinsic colonization ability of strain 11K1, but they significantly enhanced its competitive fitness (Figure 5 B). This finding indicates that increased expression of the EmhABC efflux pump improves bacterial adaptability and competitive advantage in complex rhizosphere habitats.

**Figure 5.**
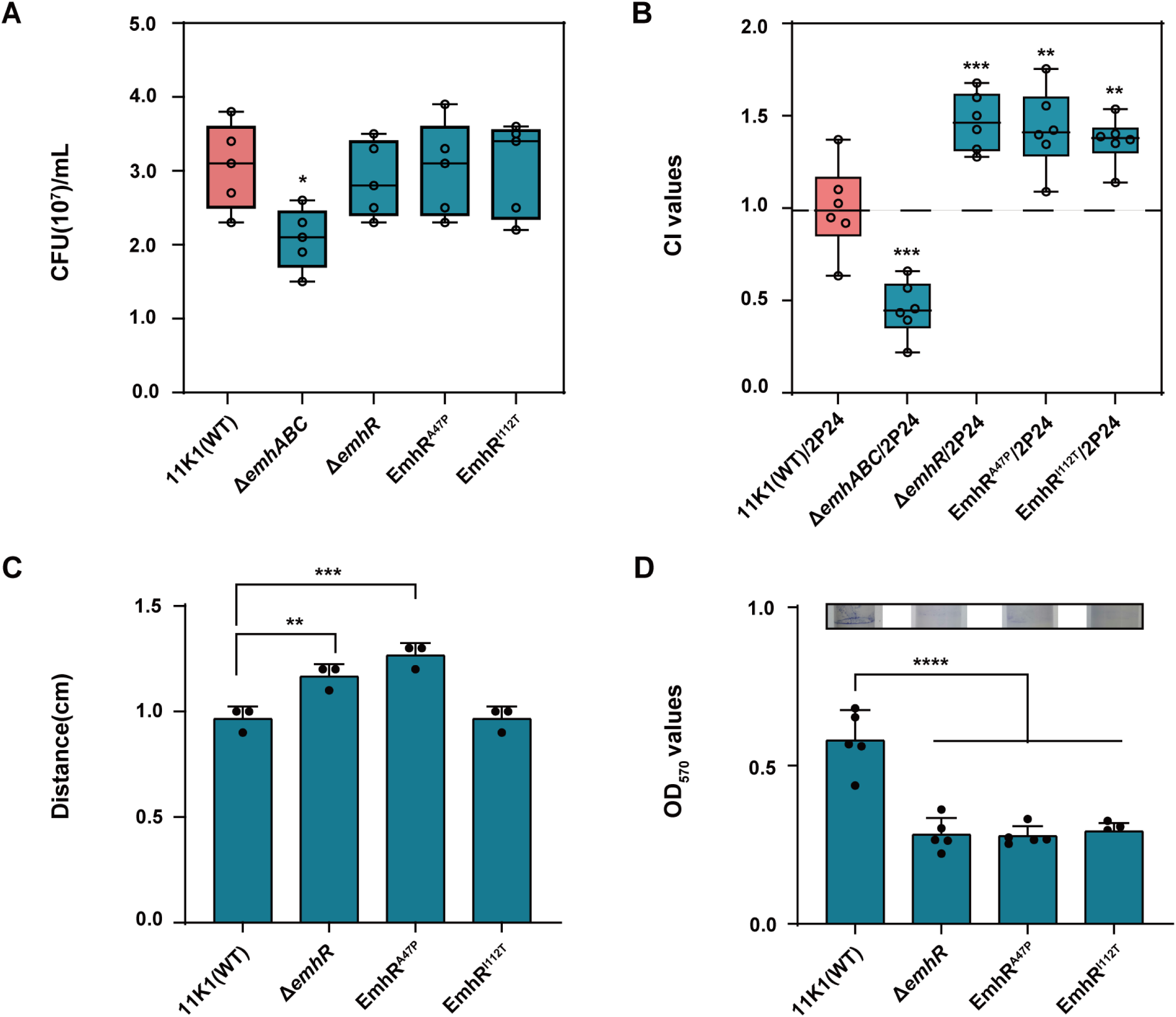
Ecological function of the A47P/I112T mutant of EmhR in the wheat rhizosphere. A. Absolute colonization levels (CFUs per gram of wheat rhizosphere soil) of 11K1, the Δ*emhR* mutant, EmhR point mutants (EmhR^A47P^, EmhR^I112T^), and the Δ*emhABC* mutant in sterilized soil. B. Competitive colonization assays of 11K1 and its derived mutants with 2P24 in the wheat rhizosphere. The competitive Index (CI) was calculated as the ratio of colony-forming units (CFUs) of 11K1 and its derivatives to those of 2P24 in the same rhizosphere niche. C. Swarming motility assays of the Δ*emhR* mutant and EmhR point mutants on swarming agar plates. D. Biofilm formation capacity of the Δ*emhR* mutant and EmhR point mutants. Statistical significance was determined by one-way ANOVA with Tukey’s post-hoc test: **P < 0.01, ****P < 0.0001.

We further evaluated the motility, biofilm formation capacity, and antimicrobial activity of EmhR mutants. Swarming assays conducted in 0.45% soft agar revealed that the Δ*emhR* and EmhR^A47P^ mutants exhibited significantly enhanced motility compared to the wild-type strain 11K1. In contrast, biofilm formation was significantly reduced in the Δ*emhR*, EmhR^A47P,^ and EmhR^I112T^ mutants relative to the wild-type (Figure 5C and 5D), indicating that EmhR mediates diverse physiological functions. Plate confrontation assays showed no significant differences in antifungal activities between the wild-type strain 11K1 and the EmhR mutants against *Thielaviopsis paradoxa* and *Colletotrichum gloeosporioides* (Supplementary Figure S4A and S4B), confirming that the EmhR mutation does not alter its antimicrobial function. Overall, the two point mutations evolved in strain 11K1 during adaptation to mupirocin, particularly EmhR^A47P^, not only confer enhanced resistance but also played critical roles in modulating bacterial motility, biofilm formation, and competitive colonizationin the wheat rhizosphere. These phenotypic changes enhance the strain’s adaptability and survival fitness in complex rhizosphere environments.

## DISCUSSION

This study utilized two rhizosphere-associated *Pseudomonas* strains to elucidate a novel mechanism of mupirocin resistance. Genetic and functional analyses revealed that the resistance is mediated by point mutations in the transcriptional regulator EmhR, which upregulates expression of the RND-type efflux pump EmhABC. This mutation-driven activation of EmhABC was identified as the primary determinant of mupirocin resistance in strain 11K1. Additionally, whole-genome sequencing analysis identified point mutations in resistant mutants within *gacA* (encoding the response regulator of the two-component regulatory system, TCS), *soxR* (a transcriptional regulator involved in oxidative stress response), and *secDF* (a heterodimeric inner membrane transporter). However, the level of mupirocin resistance conferred by these mutations was significantly lower than that associated with EmhR mutations. Numerous TCSs have been shown to regulate antibiotic resistance in Gram-negative bacteria. For instance, the VncS-VncR TCS in *Streptococcus pneumoniae* and the VanS-VanR TCS in *Enterococcus faecium* are both implicated in vancomycin resistance (39, 40). In *Acinetobacter baumannii*, the RND-type efflux pump AdeABC is regulated by the AdeR-AdeS TCS (41). In *Pseudomonas*, the GacS-GacA TCS primarily regulates the production of secondary metabolites, including antibiotics, toxins, and pigments (42, 43). As a global regulatory system, GacS-GacA can also directly or indirectly modulate the expression of various genes, including *emhABC*, thereby influencing bacterial antibiotic resistance (44, 45). In this study, *gacA* was identified as a frequently mutated gene in mupirocin-resistant isolates (Figure 3A). Nevertheless, genetic experiments demonstrated that the resistance conferred by Δ*gacA* was significantly weaker than that observed in efflux pump-related mutants (Figure 3B). Moreover, a double mutant harboring both *gacA* and *emhR* mutations was detected in one resistant isolate, 11K1-1, indicating that the *gacA* mutation is not the primary driver of mupirocin resistance in this background. Previous studies have suggested that the high mutability of *gacA* in certain *Pseudomonas* species reflects a global fitness trade-off regulatory strategy, enabling rapid metabolic reprogramming to enhance environmental adaptability (46).

The transcription regulator SoxR has been shown to primarily mediate the oxidative stress response against superoxide and nitric oxide in *E. coli* and related enteric bacteria (47, 48). However, the overexpression of SoxR in *A. baumannii* significantly reduced the expression level of the *abeS* gene, a member of small multidrug resistance (SMR) efflux pump family, by 3.8-fold, leading to markedly decreased MIC values for chloramphenicol and erythromycin. These findings indicate that SoxR functions as a negative transcriptional regulator involved in multidrug resistance in this species (49, 50). Notably, compared to RND-type efflux systems, SMR-family pumps exhibit a narrower substrate spectrum and lower transport efficiency (51). In strain 11K1, both the Δ*soxR* mutant and the L147R point mutation displayed a twofold increase in mupirocin resistance relative to the wild type (Figure 3B). Compared to the single mutations in *gacA* and soxR alone, the double mutant 11K1-2 (harboring *soxR*^L147R^ and a *gacA* frameshift mutation) exhibited significantly higher mupirocin resistance, suggesting a potential synergistic interaction between the two lesions. Nevertheless, the precise mechanism by which SoxR contributes to mupirocin resistance in 11K1 remains to be elucidated.

SecDF is a heterodimeric inner membrane transporter protein composed of the SecD and SecF subunits and classified within the RND superfamily (52). Mutations in SecDF may impair the secretion and assembly of proteins involved in bacterial drug resistance, thereby modulating bacterial susceptibility to antibiotics. In *S*. *aureus*, loss of SecDF function disrupts cell wall homeostasis, resulting in increased resistance to β-lactam antibiotics and glycopeptides (53). In this study, the individual absence of either SecD or SecF did not affect the resistance of strain11K1 to mupirocin (Figure 3B). However, in 11K1-52, which harbors the mutations SecD^D100E^, SecF^P120S^, and GacA^L27R^ (Figure 3A), the observed enhancement in mupirocin resistance may result from the synergistic effects of GacA and SecDF mutations, or may involve additional mechanisms that require further elucidation.

The EmhR-EmhABC efflux system was first identified in *P. fluorescens* and belongs to the RND efflux pump family (54). EmhR functions as a transcriptional repressor located upstream of the *emhABC* gene cluster where it negatively regulates the expression of this operon, thereby contributing to bacterial resistance to multiple antibiotics and toxic ions (36, 55). In this study, two point mutations in the EmhR protein were found to significantly enhanced the resistance of strain 11K1 to mupirocin. Notably, the residue A47 is positioned in the α3 helix of EmhR, adjacent to the DNA-binding interface of the EmhR dimer (Figure 3D). EMSA assays further demonstrated that the mutation at this site significantly impairs the binding affinity between EmhR and its target DNA sequence (Figure 3E). A similar regulatory mechanisms has been reported for MexL, a paralog of EmhR in *P. aeruginosa*. MexL acts as a transcriptional regulator for the RND efflux pump MexJK-OprM. The MexL^A47D^ mutation (corresponding to A41 in EmhR of 11K1) introduces a hydrogen bond between D47 and R29 in helix 1, which severely hinders MexL binding to the mexJ promoter region, ultimately leading to derepression and increased expression of the efflux pump (56). In this study, the A47P substitution in 11K1 emerged spontaneously under mupirocin selection pressure and was independently observed in multiple mutant strains (11K1-1 and 11K1-6) (Figure 3A), suggesting that this mutation confers a significant selective advantage by enhancing bacterial survival and adaptability in the presence of mupirocin. In another mutant, the I112 is situated near the dimerization interface of EmhR. Structural studies have shown that nearly all resolved TetR-family proteins, including EmhR, adopt a homodimeric configuration (57, 58), which optimally positions the helix-turn-helix (HTH) domain for specific DNA base recognition (59). Mutations disrupting dimer formation may compromise the structural integrity required for precise DNA binding, thereby altering the regulation of downstream genes (60). In comparson, the A47P mutation demonstrates a more pronounced effect on modulating EmhR function, as further supported by EMSA and RT-qPCR data (Figure 3E and Figure 4A).

Currently, the primary efflux pump families associated with bacterial multidrug resistance include the RND family, the major facilitator superfamily (MFS) family, the multidrug and toxic compound extrusion (MATE) family, and the SMR family proteins (61). A common structural feature of these efflux pumps is the presence of transmembrane domains, which enable the recognition and binding of diverse substrates and facilitate their energy-dependent expulsion from the cell, thereby conferring a drug-resistant phenotype to the bacteria. In contrast to the MFS, MATE, and SMR family efflux pumps, which are primarily localized to the inner membrane system and exhibit relatively simple architecture, the multi-component RND efflux pump system can directly transport substrates from the cytoplasm or inner membrane to the extracellular space. This capability makes them the most clinically significant class of efflux pumps associated with drug resistance in Gram-negative bacteria (62, 63). Besides the overexpression of efflux pump genes, which directly enhances antimicrobial resistance, mutations in efflux pump structural genes, as well as the integration of mobile genetic elements such as integrons, leading to the constitutive expression of these pumps, can significantly increase bacterial resistance to antibiotic (64, 65). Furthermore, efflux pump-mediated resistance typically arises through the synergistic interplay of multiple mechanisms, including target site modification and enzymatic inactivation of antimicrobial agents (66). Importantly, efflux pump systems are not solely involved in antimicrobial resistance but also contribute to other critical biological functions, such as secretion of quorum-sensing signaling molecules and bioactive compounds (e.g., 2,4-DAPG), biofilm formation and the export of virulence factors (36, 67, 68). These multifaceted roles underscore the importance of efflux pumps in bacterial survival and adaptability, explaining their conservation and evolutionary persistenceeven in antibiotic-free environments.

Bacterial antibiotic resistance is not merely a survival response to antimicrobial pressure but also serves as a crucial mechanism facilitating colonization and competitive fitness within complex ecological niches. The RND efflux pump system has been shown to be essential for *Vibrio cholerae* in establishing antibiotic resistance, optimizing virulence factor expression, and enabling intestinal colonization in infant mice (69). Similarly, the AcrAB-TolC-mediated efflux of bile salts plays a critical role in allowing *E. coli* to successfully colonize the human intestinal environment. Notably, most such studies have extensively focused on clinical pathogens, with considerably fewer investigations into analogous mechanisms in environmentally derived strains (70, 71). In this study, two point mutations in EmhR did not alter the absolute colonization levels of strain11K1 in the rhizosphere; however, compared to the wild-type 11K1, these mutations significantly enhanced the competitive colonization ability of 11K1 against strain 2P24 in the wheat rhizosphere (Figure 5A, Supplementary Figure S5A). This suggests that the mutations markedly improved the ecological adaptability of 11K1 within the rhizosphere community. Vancomycin-resistant *enterococci* (VRE) can gain a significant competitive advantage in the antibiotic-treated gut (72), and the resistance acquisition occurs not only through intrinsic genetic mutations but also via horizontal gene transfer mechanisms, including plasmids, transposons, and the capture of resistance genes via integrons (73, 74). Clinically, mupirocin is used in the treatment of methicillin-resistant *Staphylococcus aureus* (MRSA). As a natural producer of mupirocin, *Pseudomonas* spp., together with its specialized ecological niche in the rhizosphere, provides valuable insights into the evolutionary origins and mechanisms of mupirocin resistance. Rhizosphere-associated bacteria may have inherently evolved resistance mechanisms, which are of considerable significance for understanding the natural reservoirs of antimicrobial resistance and for informing the development of novel therapeutic strategies in clinical practice.

## MATERIALS AND METHODS

### Microbial strains and growth conditions

The bacterial strains and plasmids used in this study are detailed in Table S1. *P. bijieensis* 2P24 (2P24), *P. viciae* 11K1 (11K1), and their derivative strains were grown in LB medium, King’s B (KB) medium (75) or Potato Dextrose Agar (PDA) medium at 28°C. *P. aeruginosa* PAO1 (PAO1) and its derivatives were cultured in LB broth or AB minimal (ABM) medium (76) at 30°C, whereas *E. coli* BL21(DE3) and DH5α were cultured in LB broth at 37°C. The pathogenic fungi *Thielaviopsis paradoxa* and *Colletotrichum gloeosporioides* were cultured on PDA medium at 30°C. When required, antibiotics were added at the following concentrations: Kanamycin (Km), 50 µg/mL; Gentamicin (Gm), 30 µg/mL; Tetracycline (Tc), 20 µg/mL; and Ampicillin (Amp), 50 µg/mL.

### Plate confrontation assays

Strain 2P24 and its derivatives were cultivated in 30 mL tubes containing 3 mL of LB broth with shaking at 200 rpm for 16 h at 28°C. Bacterial cells were harvested by centrifugation, resuspended, and adjusted to an optical density (OD_600_) of 0.5 (~1×10^8^ cells mL^−1^) in sterile water to prepare a bacterial suspension. A 5 μl aliquot of this suspension was placed at the center of a PDA plate and incubated statically for 16 h at 28°C. Following the same protocol, a suspension of strain 11K1 was prepared, and 5 μl aliquots were spotted at distances of 10 mm and 20 mm from the 2P24 colony. Plates were allowed to dry under ambient conditions before being incubated statically for 24 h at 28°C. All experiments were performed with at least three biological replicates and three technical replicates.

### Screening and generation of mupirocin-resistant mutants of 11K1

Prepare the 11K1 bacterial suspension (1×10^8^ cells mL^−1^) as previously described. Subsequently, mix 200 μL of this suspension into molten KB agar medium prior to pouring plate. A 10 μL volume of a 10 mg/mL mupirocin solution was applied at the center of each plate, and an equivalent volume of DMSO was added in control plates. The Petri dishes were sealed and incubated statically at 28°C for 48 h. The mutants were picked and streaked aseptically for single-colony isolation. Minimum inhibitory concentration (MIC) assays are performed to evaluate mupirocin resistance in these isolates. Isolates exhibiting significantly higher resistance compared to wild-type 11K1 are designated as 11K1 mupirocin-resistant mutants.

### MIC Assay

The MIC assay was performed using the standard 96-well plate microdilution method (77). Briefly, Strain 11K1 and its derivatives were cultured overnight at 28°C with shaking at 200 rpm in LB broth. Bacterial suspensions were adjusted to an OD_600_ of 1.0 (~1×10^9^ cells mL^−1^) in sterile water and subsequently diluted 1:10,000 in fresh LB broth. A volume of 200 μL of the diluted inoculum (~1×10^5^ cells mL^−1^) was dispensed into each well of a sterile polystyrene 96-well plate. mupirocin or other antibiotics were added to the wells in two-fold serial dilutions across the wells, with corresponding solvent controls included for each compound. The plates were incubated statically at 28°C for 24 h. Following the incubation, bacterial growth was quantified by measuring the OD_600_ using a multimode microplate reader (SpectraMax i3x), and the MIC was determined as the lowest antibiotic concentration that completely inhibited bacterial growth.

### Single nucleotide polymorphisms (SNPs) detection

Genomic DNA was extracted from the parental strain 11K1 (designated as the starting strain in this study) and the 11K1 mupirocin-resistant mutant strain using the CTAB method. SNP detection was carried out according to previously established protocols (78). Briefly, sequencing libraries were prepared using the NEBNext® Ultra™ DNA Library Prep Kit for Illumina. Following confirmation of library quality, pair-end sequencing (PE150) was performed on Illumina NovaSeq platform to achieve a sequencing depth of approximately 200-400x for each sample. Low-quality reads (defined as those containing more than 50% of bases with a quality score ≤5) were filtered out prior to analysis. The resulting high-quality reads were then aligned to the reference genome using BWA (v0.7.17). SNP calling for individual strains was subsequently conducted using GATK (v4.0.5.1).

### Construction of 11K1 and PAO1 mutants and complementation strains

The 11K1 deletion and point mutation strains were generated using a two-step homologous recombination strategy. All primers used in this study are listed in Table S2. As a representative example, the upstream and downstream fragments of the *emhR* gene in strain 11K1 were amplified by PCR using primer pairs *emhR*-UF/UR and *emhR*-DF/DR, respectively, and purified prior to cloning. These fragments were then cloned into the plasmid p2P24Km (79) to construct the suicide vector p2P24-*emhR*. The resulting plasmid was validated by Sanger sequencing before being introduced into strain 11K1 via electroporation. Putative transformants were initially selected in LB medium containing Km (50 μg/mL), followed by counter-selection on LB medium supplemented with 15% sucrose. The Δ*emhR* deletion mutant was ultimately confirmed by PCR amplification and Sanger sequencing. Similarly, the single-nucleotide mutant strains were conducted using a procedure analogous to that described above.

The construction of the PAO1 deletion mutant strains were performed following previously reported methods with minor modifications (80). The two-step homologous recombination technique was employed, using the suicide plasmid pEX18 (Gm) (56). The *E. coli* S17-1 λpir strain, harboring the recombinant suicide plasmid, was used as the donor in conjugation experiment with PAO1 to enable the transfer of the plasmid into the recipient strain. Subsequent screening procedures were analogous to those employed in the construction of the 11K1 mutants. For complementation, the *nalD* gene was amplified by PCR and cloned into the shuttle plasmid pBBR1MCS-5, yielding the recombinant plasmid pNalD. This plasmid was verified by Sanger sequencing and subsequently introduced into the *nalD* deletion mutant. The same approach was applied to complement the NalD point mutation.

### Expression and purification of His6-EmhR Protein

The *emhR* gene fragment was amplified via PCR from the 11K1 genomic DNA using specific primers R6His-F/R (Table S2). The purified PCR product was ligated into the linearized pET22b vector to generate the recombinant protein expression vector pET22b-EmhR. Using the same approach, the corresponding expression vectors pET22b-EmhR^A47P^ and pET22b-EmhR^I112T^ were constructed. After verification by Sanger sequencing, each plasmid was introduced into *E. coli* BL21 (DE3) for protein expression. Transformed cells were cultured in LB broth containing Km (50 μg/mL) at 37°C with shaking at 150 rpm until the OD600 reached 0.6. The culture was then shifted to 16°C and induced with 0.25 mM IPTG for 16 h to promote the target protein expression. Protein purification was performed according to a previously described protocol (33), and the final concentration of the purified protein was determined to be 4-5 mg/mL.

### RNA extraction and qRT-PCR analysis

Total RNA was extracted from bacterial cells as described previously with minor modifications (79). Briefly, bacterial cell suspensions were collected at appropriate time points and centrifuged at 13600 × g for 2min to harvest the pellets. Cells were lysed using Trizol kits (Invitrogen, Shanghai, China), followed by phase sparation with chloroform to remove proteins and other organic components. Residual DNA was eliminated using FastKing gDNA Dispelling RT SuperMix (Tiangen Biotech, Beijing, China), and first-strand cDNA was synthesized from total RNA via reverse transcription. Quantitative reverse transcription PCR (qRT-PCR) was performed on a real-time fluorescent quantitative PCR instrument (JLM, Sichuan, China) using a 96-well plate format. The 20 μL reaction mixture contained 10 μL of SYBR Green Mix (TaKaRa, Dalian, China), 1 μLof cDNA template (500 ng), and 2 μL of gene-specific primers. Amplification was carried out under the following conditions: initial denaturation at 95°C for 30 seconds, followed by 40 cycles of 95°C for 5 seconds and 60°C for 30 seconds. The 16S rRNA gene was used as an internal reference for normalization of target gene expression levels. Relative changes in gene transcription were calculated using 2^−ΔΔCt^ method, where ΔCt = Ct_gene of interest_ −Ct_16S rRNA_ and Ct denotes the threshold cycle.

### Electrophoretic mobility shift assay (EMSA)

The nucleotide sequence of the EmhABC promoter region (255 bp) was amplified from the 11K1 genome by PCR using the primers p*emhABC*-F/R. The interaction between EmhR and p*emhABC* was evaluated by electrophoretic mobility shift assay (EMSA). In brief, the p*emhABC* probe was incubated with varying concentrations of purified recombinant proteins His6-EmhR, His6-EmhR^A47P^, and His6-EmhR^I112T^ in a DNA binding buffer (20 mM Tris-HCl, pH 8.0, 20 mM NaCl, and 10% glycerol) at room temperature for 30 minutes. Following incubation, the reaction mixtures were resolved on a 12% non-denaturing polyacrylamide gel in electrophoresis buffer (89 mM Tris-borate, pH 8.0), 2 mM EDTA) at 100 V under cold conditions. The gel was subsequently visualized and digitally captured using the ImageQuant TL10 imaging system.

### Competitive colonization assays

The mini-Tn7 site-specific insertion method (81) was employed to introduced plasmids pCPP6529 (Km) and pCPP6351 (Tc) (82) into the wild-type strains 2P24 and 11K1, as well as their respective mutant derivatives, for antibiotic resistance labeling. Single colonies confirmed by Sanger sequencing were inoculated in LB broth and incubated at 28°C with shaking at 200 rpm for 16 hours. Bacterial cells were harvested by centrifugation and resuspended in sterile water to a final OD_600_ of 0.5 (approximately 1×10^8^ cells mL^−1^). Healthy germinating wheat seeds were soaked in a 1:1 (v/v) mixture of the 2P24 and 11K1 bacterial suspensions for 30 minutes. After air-dring, the seeds were sown in a sterile mixture of peat and vermiculite, which had been autoclaved twice (121°C, 20min). Two weeks post-planting, root tissues were collected with approximately 1 mm of adhering substrate. The root samples were washed in sterile water, and the resulting suspensions were serially diluted and plated on selective antibiotic media for colony enumeration, followed by statistical analysis.

## SUPPLEMENTAL MATERIAL

ASM does not own the copyrights to Supplemental Material that may be linked to, or accessed through, an article. The authors have granted ASM a non-exclusive, world-wide license to publish the Supplemental Material files. Please contact the corresponding author directly for reuse.

## ACKNOWLEDGEMENTS

This work was funded by the National Key Research and Development Program of China (2023YFA0915900, 2025YFC2608200) and the National Natural Science Foundation of China (32372497, 32272611).

We sincerely acknowledge the Laboratory of Agricultural and Forestry Biosecurity (MARA Key Lab of Pest Monitoring and Green Management), and the Public Experimental Platform of the College of Plant Protection, China Agricultural University for their assistance and support in this study.

## Supplementary Figure Legends

Supplementary Figure S1 Experimental design of mupirocin MIC assays and biological characteristics of strain 11K1 and its mupirocin-resistant mutants. A. Schematic representation of the mupirocin MIC assay performed via microdilution in LB medium. B–C. Growth curves of 11K1 and mupirocin-resistant mutants in LB medium (B) and M9 minimal medium (C), monitored by measuring OD₆₀₀ values at indicated time points under shaking conditions (at 28°C, 200 rpm). D. Colony morphology of 11K1 and its mupirocin-resistant mutants following single-colony streaking on PDA plates.

Supplementary Figure S2 Specificity of EmhR binding to the promoter p*emhABC*. FAM-labeled EmhABC promoter DNA (FAM-p*emhABC*) was incubated with His₆-EmhR in the presence of increasing concentration of unlabeled p*emhABC*. The binding of FAM-p*emhABC* to EmhR was competitively inhibited by unlabeled p*emhABC* in a concentration-dependent manner, confirming the specificity of the EmhR-DNA interaction.

Supplementary Figure S3 Transcriptional regulation of *emhABC* by EmhR deletion and point mutations. Transcriptional expression levels of *emhA*, *emhB*, and *emhC* in strain 11K1, the Δ*emhR* mutant, and EmhR point mutants (EmhR^A47P^, EmhR^I112T^) at 18 hpi, quantified by qRT-PCR. Statistical significance was determined by one-way ANOVA with Tukey’s post-hoc test: **P < 0.01, ****P < 0.0001.

Supplementary Figure S4 Plate confrontation assays of the Δ*emhR* mutant and EmhR point mutants against the fungal pathogen *Colletotrichum gloeosporioides* (A) and *Thielaviopsis paradoxa* (B). Inhibition zones were observed after 48 h of co-incubation at 28°C.

